# Group Size as a Trade Off Between Fertility and Predation Risk: Implications for Social Evolution

**DOI:** 10.1101/342550

**Authors:** R.I.M. Dunbar, Padraig Mac Carron

## Abstract

Cluster analysis reveals a fractal pattern in the sizes of baboon groups, with peaks at ∼20, ∼40, ∼80 and ∼160. Although all baboon species individually exhibit this pattern, the two largest are mainly characteristic of the hamadryas and gelada. We suggest that these constitute three pairs of linear oscillators (20/40, 40/80 and 80/160), where in each case the higher value is set by limits on female fertility and the lower by predation risk. The lower pair of oscillators form an ESS in woodland baboons, with choice of oscillator being determined by local predation risk. Female fertility rates would naturally prevent baboons from achieving the highest oscillator with any regularity; nonetheless, hamadryas and gelada have been able to break through this fertility ‘glass ceiling’ and we suggest that they have been able to do so by using substructuring (based partly on using males as ‘hired guns’). This seems to have allowed them to increase group size significantly so as to occupy higher predation risk habitats (thereby creating the upper oscillator).

## Introduction

Groups are of fundamental importance in an evolutionary context for two reasons. First, they provide the demographic context in which animals play out their individual social and reproductive strategies. The size and demographic structure of the group in which an individual lives determines its behavioural and strategic options: if you don’t have a sister, you cannot form an alliance with sisters, even though this may be the optimal thing to do (Altmann & Altmann 1979; Dunbar 1979). Second, group size is the key interface between the animal and its environment and, because primates, in particular, solve many of their ecological challenges socially, group size will often be a crucial feature of their ecological success. Understanding the factors that determine the size of groups that individuals live in is thus essential if we are to fully understand the processes that lie behind primate evolutionary history, and the particular behaviour of extant species. Despite this, the determinants of group size for any primate species have not been a significant focus of interest for several decades, and our understanding remains disappointingly poor and often very informal.

For all animals, group size is ultimately a tradeoff between costs and benefits. For primates (and most cursorial mammals), the principal benefits arise from protection against predation (Altmann & Altmann 1970; Cowlishaw 1994; Shultz et al. 2004; Cheney et al. 2004; Adamczak & Dunbar 2008; Lehmann & Dunbar 2009a; Willems & Hill 2009; Shultz & Finlayson 2010; Bettridge & Dunbar 2012). The costs arise from a combination of indirect costs (increased day journey length and foraging demand: Dunbar et al. 2009) and direct costs (competition between group members, and in particular the consequences this has for female fertility: Dunbar 2018). In this paper, we explore how these play out to influence social group size in baboons.

Predation by cursorial predators (principally lion, leopard and hyaena) is a serious problem faced by baboons (Altmann & Altmann 1970; Cowlishaw 1994; Bettridge & Dunbar 2012), especially at night when these predators are most active and primates are vulnerable due to their poor night vision. Not only do the densities of predators place constraints on the biogeographic distributions of all terrestrial primates (Lehmann & Dunbar 2009a), they also dictate minimum group size (Dunbar et al. 2009; Bettridge & Dunbar 2012). Some evidence of the significance of this for baboons is provided by a rare case of group fusion: two groups in the Mikumi National Park (Tanzania) whose sizes had progressively declined over time to 11 and 13 due to high mortality rates fused to create a group of 24 individuals (Hawkins 1999). Not only were both of the original groups below the size for demographic viability (∼17 individuals: Dunbar et al. 2018), but they were also below the minimum group size (16.3) predicted for this site by the local predator density (Bettridge et al. 2010). Fusing placed both groups in the demographic ‘safe zone’.

In contrast, the substantive costs of group-living seem to arise from the impact that this has on female fertility, rather than access to resources *per se.* In mammals, female reproductive endocrinology is extremely sensitive to stress: even low levels of stress can destabilise it, resulting in increased levels of infertility, and in some cases even the complete suppression of puberty (Bowman et al. 1978; Abbott et al. 1981, 1986, 1988; Seifer & Collins 1990). The mechanism for this is well understood pharmacologically: high levels of ß-endorphins produced in response to social or physical stress block the production of GnRH in the hypothalamus, with the result that the pituitary fails to produce the lutenising hormone (LH) surge required to trigger ovulation, leading to anovulatory (and hence infertile) menstrual cycles (Zacur et al. 1976; Seifer & Collins 1990; Laatikainen 1991; von Borell et al. 2007; Einarsson et al. 2008). A decline in fertility with either rank (Dunbar 1980; Smuts & Nicholson 1989; Altmann & Alberts 2003; Garcia et al. 2006; Pusey & Schroepfer-Walker 2013) or group size (van Schaik 1983; Dunbar 1989; Srivastava & Dunbar 1996; Hill et al. 2000; Borries et al. 2008) has been widely documented in primates, and seems to be characteristic of mammals as a whole (Dunbar 2018). Since the same endocrinological mechanism underpins human and primate infertility (An et al. 2013; Schliep et al. 2015; Pettay et al. 2016), and stress has been an identified factor in human infertility (An et al. 2013; Schliep et al. 2015; Pettay et al. 2016), similar effects have also been noted in humans: fertility declines with harem size in ethnographic societies (Muhsam 1956; Smith & Kunz 1976; Chojnacka 1980; Bean & Mineau 1986; Borgerhof Mulder 1989).

Ecologists have invariably attributed the decline in fertility with rank or group size to reduced food intake as a result of scramble competition. In fact, starvation itself triggers the endorphin system and thereby precipitates infertility (Dobson et al. 2012, Clarke 2014), so the effect may be due to stress rather than shortage of nutrients *per se*. While lack of food can certainly cause the reproductive system to shut down, this usually happens only in cases where extreme starvation or excessive exercise (e.g. athletes) results in insufficient fat reserves to support pregnancy (Smith 1947; Dean 1949; McClure 1968; Warren & Perlroth 2001; Kirchengast & Huber 2001). Even then, it seems to be the hypothalamic pathway that regulates this, rather than nutrition itself (Kalra & Kalra 1996; Schwartz & Seeley 1997). Indeed, experimental studies of domestic stock have shown that stocking rate (in effect, social group size) is responsible for this, independently of any effect due to food availability or nutrition (von Borell et al. 2007; Einarsson et al. 2008; Dobson et al. 2012; Clarke 2014). In short, it seems that the same endorphin/HPA pathway is involved in both social and ecological routes, perhaps explaining why it can be difficult to distinguish between their respective effects. In short, it seems that living in large groups can adversely affect female fertility irrespective of how rich the foraging conditions may be.

For species that live in loose associations (herds), these costs can easily be defused by leaving the group and joining a smaller one – as is widely exemplified in non-primate mammals. However, for species like primates that live in bonded social groups (Dunbar & Shultz 2010; Shultz & Dunbar 2010; Massen et al. 2010), individuals cannot easily leave one group to join another because entry by strangers is often aggressively resisted (Dunbar 1984; Kahlenberg et al. 2008). Since predation risk militates against leaving groups to forage alone (Dunbar et al. 2009), the only way a group can lose members is to fission into two independent groups large enough to survive on their own. Fission has been documented in many primate species (e.g. Dunbar 1988; Ménard & Vallet 1993; Henzi et al. 1997; van Horn et al. 2007). An important constraint is that if minimum group size is set by the local predation risk (Dunbar et al. 2009; Bettridge & Dunbar 2012), groups need to be at least twice this size at fission in order to allow both daughter groups to be above the specified minimum (see also Dunbar & Sosis 2017).

We use a comprehensive dataset of baboon group sizes to investigate the structural patterns that these exhibit, and explore likely reasons for the patterns in terms of predation risk and fertility. We take baboons to refer to the large terrestrial primates of Africa (‘baboons’ *sensu* Jolly 2007). For present purposes, we include the genera *Papio* (the baboons of conventional usage) and *Theropithecus* (the gelada). We exclude *Mandrillus* (drills and mandrills) because their groups are still poorly understood and have yet to be reliably censused. There are some grounds for considering *Lophocebus* and *Rungwecebus* as allies of the baboons, but as these are arboreal and hence likely to be subject to very different selection pressures as a result (arboreal habitats are typically much less predator risky), we exclude them as well.

We ask three questions: (1) Is there a typical or natural size for baboon groups? (2) Why do baboons have the range of group sizes that they do? and (3) How can we explain the structural differences that distinguish some species (specifically, hamadryas and gelada with their multilevel harem-based social systems) from others (the remaining four species of *Papio* which lack a harem structure).

## Methods

We comprehensively searched the literature for census data on baboon group sizes, providing such censuses were part of a systematic attempt to count the size and composition of individual baboon groups at a given location. We take group (or troop) size to be that defined by the field worker. For *Papio anubis*, *P. cynocephalus* and *P. ursinus*, the group is fairly obvious, since animals forage and sleep together in stable social groups that maintain a degree of demographic coherence and stability over time as well as spatial separation from neighbouring groups. Exactly what counts as a group in *Papio papio* has been the subject of some debate because of this species’ rather more flexible form of sociality, but all field workers on this species agree on the existence of some form of stable social group, at least as defined by animals that share a common range area (Dunbar & Nathan 1972; Boese 1975; Sharman 1981; Patzelt et al. 2011). For *P. hamadryas* and *Theropithecus gelada*, we equate the social group with the band (a collection of single male reproductive units, or harems, that share a common home range and frequently forage together: Kummer 1968; Mac Carron & Dunbar 2016). In all three latter cases, we use the group (or band) as defined by the fieldworker concerned. However, in order to ensure that we were comparing like with like, we excluded publications (e.g. Zinner et al. 2001) which provided only mean group sizes for a location or did not provide clear evidence for differentiating between grouping levels in the case of those species (*P. hamadryas* and *T. gelada*) which have multilevel societies.

In populations that were subject to longterm study with repeated censuses of the same social groups, we counted a census as being new only providing it had been carried out at least five years after the previous census of that group. In either case, we always used the original (i.e. earliest) census and at most one later census. This yielded a total of 444 groups across 53 study sites in 13 countries for five species of *Papio* and the one extant species of *Theropithecus*. There are insufficient data available for *P. kindae*, which has only recently been elevated to species status (Zinner et al. 2013) and has not been well studied in the field. The data are given in Table S1.

We first determined whether the distribution of group sizes is unimodal. If the distribution is not normal, our task is to find the most appropriate distribution that does describe the data. We used maximum-likelihood methods (Clauset et al. 2009) to fit the following common distributions: power law, truncated power law, geometric, negative binomial, exponential, stretched exponential, normal, lognormal, and a compound Poisson distribution. The first eight of these assume that the data are non-normal (i.e. skewed) distribution with a single definable mode. The last assumes that the observed distribution is in fact the product of several separate distributions, each with their own modes, that have been pooled together. That might occur because each species has a single modal (or typical) value, but the values for the various species differ significantly. However, more interestingly, such a pattern might occur when a group’s size oscillates within sets of defined limits, with the particular limits being set by some key aspect of the animals’ ecology; in these cases, individual species may be found in a wide range of group sizes (or group size “types”).

As the dataset consists of discrete integers, we used the discrete approach as described by Clauset et al. (2009). We numerically maximised the log-likelihood of each candidate distribution to obtain its parameter estimates, using the *optimize* module of Python’s *scipy* (v0.17.1) library, and identified the most likely model using the Akaike Information Criterion (AIC). In the case of the compound Poisson, we treated the data as being made up of 1 to *n* Poisson distributions; the AIC for each *n* is calculated successively, stopping when a local minimum is reached to give the optimal *n*. This is the only distribution tested that is multimodal; the rest have a single peak and, except for the normal distribution, all are right skewed.

Because models with more parameters are always more likely to fit the data, we applied a clustering algorithm to see if a different approach gives the same result. Since most clustering algorithms are meant for a high number of dimensions (Jain 2010), we used the Jenks natural breaks algorithm as this is more appropriate for data with only one dimension (Jenks 1967). Jenks is very similar to *k*-means clustering in one dimension and works by minimising the variance within different numbers of clusters. In order to choose the optimal number of clusters, we calculated a goodness of fit for each number of clusters. Since a goodness of fit of 1.0 can only be attained when there is no within-class variation (typically when the number of clusters is the same as the sample size), we follow Coulson (1987) and take a value of 0.85 as the threshold.

Female fertility rates for 15 individual baboon troops are taken from Hill et al. (2000), with additional data from more recent studies for one group each for *P. anubis* (Higham et al. 2009) and *P. ursinus* (Cheney et al. 2004), two groups for *P. hamadryas* (Swedell & Saunders 2006; Chowdhury et al. 2015), and three bands for *T. gelada* (from Ohsawa & Dunbar 1984). In addition, we give two population means for *P. papio* (Boese 1975; Sharman 1981) based on the number of immatures (prepuberty individuals, less than ∼ 4 years of age) in a group. In plotting fertility rate against group size, we use the mean size of the foraging group in the case of the multilevel hamadryas and gelada since this is the social context in which animals spend most of their time and it is this that affects female fertility. For hamadryas, we identify this as the band; for gelada, we identify this as the herd. Mean population group size, climatic and forest cover data for these habitats are from Bettridge et al. (2010). The data are provided in Table S2.

## Results

### Group size

The distribution of baboon group sizes is highly skewed, with a mean of 48.5±43.4SD and a range of 3-262 (Fig. 1). Analysis of variance indicates that group sizes vary significantly across the six species (Fig. 2: F_5,438_=32.25, p<<0.0001). Scheffé post hoc tests suggest that the data fall into two clusters that differ significantly from each other (p<0.001). These are: (1) *Papio anubis* (mean group size = 40.3±26.4, N=104), *P. cynocephalus* (mean group size = 51.7±34.5, N=105) and *P. ursinus* (mean group size = 30.4±21.9, N=166) and (2) *P. papio* (mean group size = 91.6±69.4, N=34), *P. hamadryas* (mean group size = 93.5±62.4, N=13) and *Theropithecus gelada* (mean = 115.7±71.7, N=22). For convenience, we label the first set as the ‘woodland’ species, a term that has no taxonomic implications but refers to the fact that they mostly occupy savannah woodland habitats that are rather different to the kinds of habitats occupied by the hamadryas and gelada.

**Figure 1.**
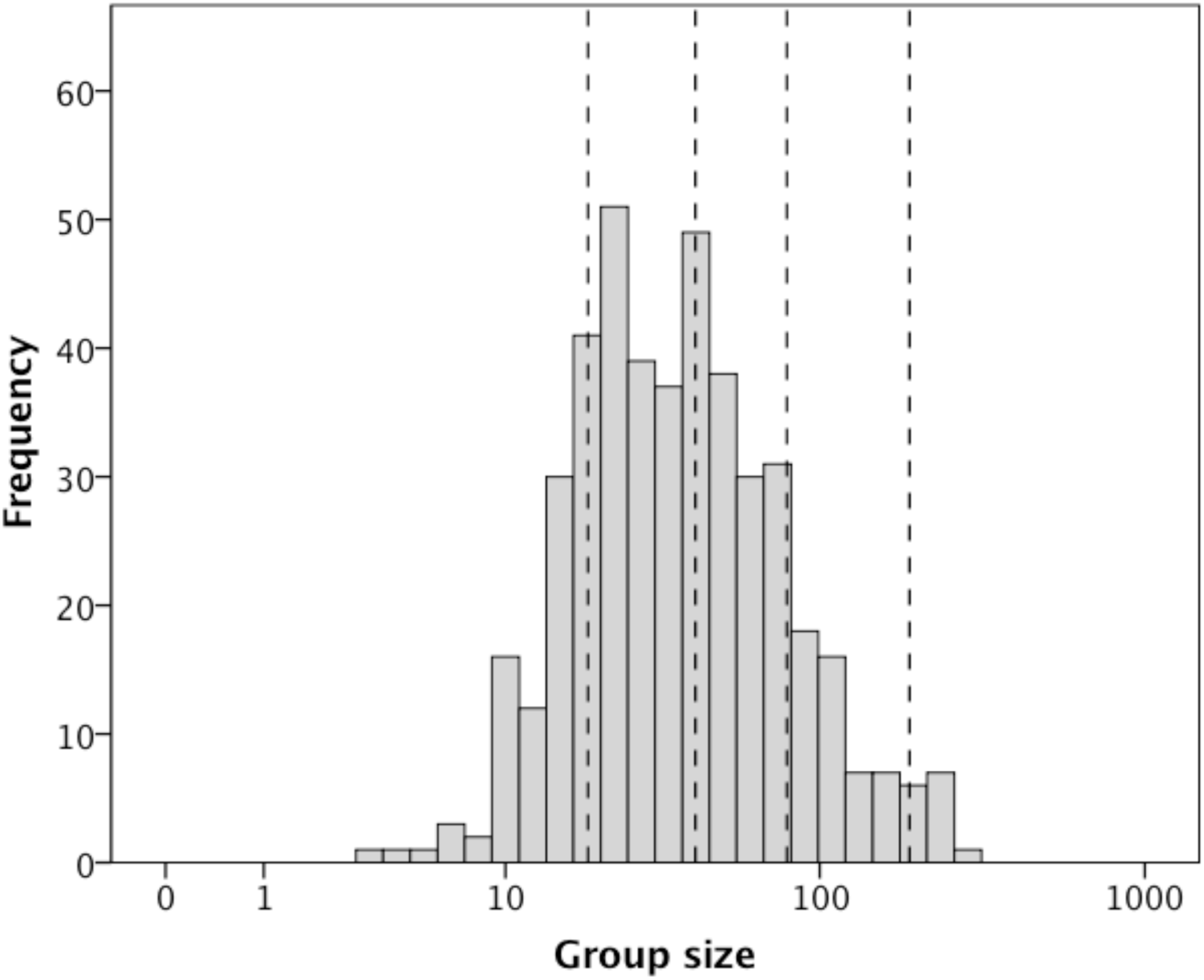
Distribution of social group sizes in baboon (*Papio* and *Theropithecus*) populations. Dashed vertical lines demarcate the optimal group sizes of 18.7, 41.0, 79.1 and 180.3, averaged for the two the maximum likelihood estimates (see text for details).

**Figure 2.**
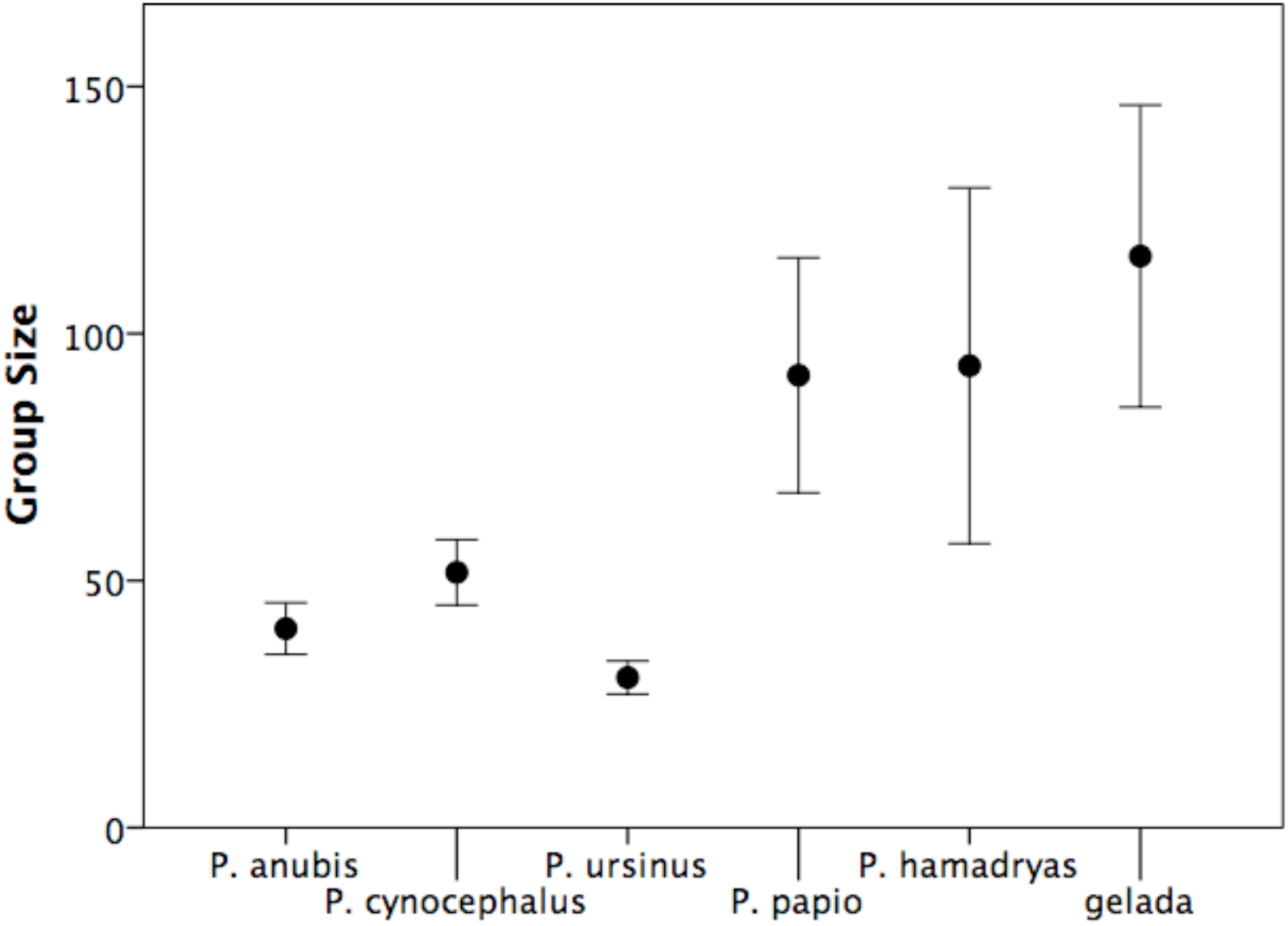
Mean (±2 se) group size for individual species. Samples size are: *P. anubis* = 102, *P. cynocephalus* = 109, *P. ursinus* = 165, *P. hamadryas* = 13; *P. papio* = 34 groups; *Theropithecus gelada* = 22 groups.

Applying maximum likelihood estimation to the raw data in Fig. 1 indicates that the distribution is most likely made up of 4 Poisson distributions (Table 1) with means 19.1, 42.6, 83.3 and 189.4. Excluding gelada gives virtually the same results (means at 18.7, 41.0, 79.1 and 180.3). The Jenks algorithm also suggests that four clusters is optimal for all *Papio* combined (means of 19.8, 45.6, 85.4 and 184.4), but opts for just three clusters when gelada are included (means 27.1, 77.0 and 190.1). All four series have a mean scaling ratio of ∼2.1 (2.7 for the last case), suggesting a fractal pattern indicative of binary fission.

**Table 1.**
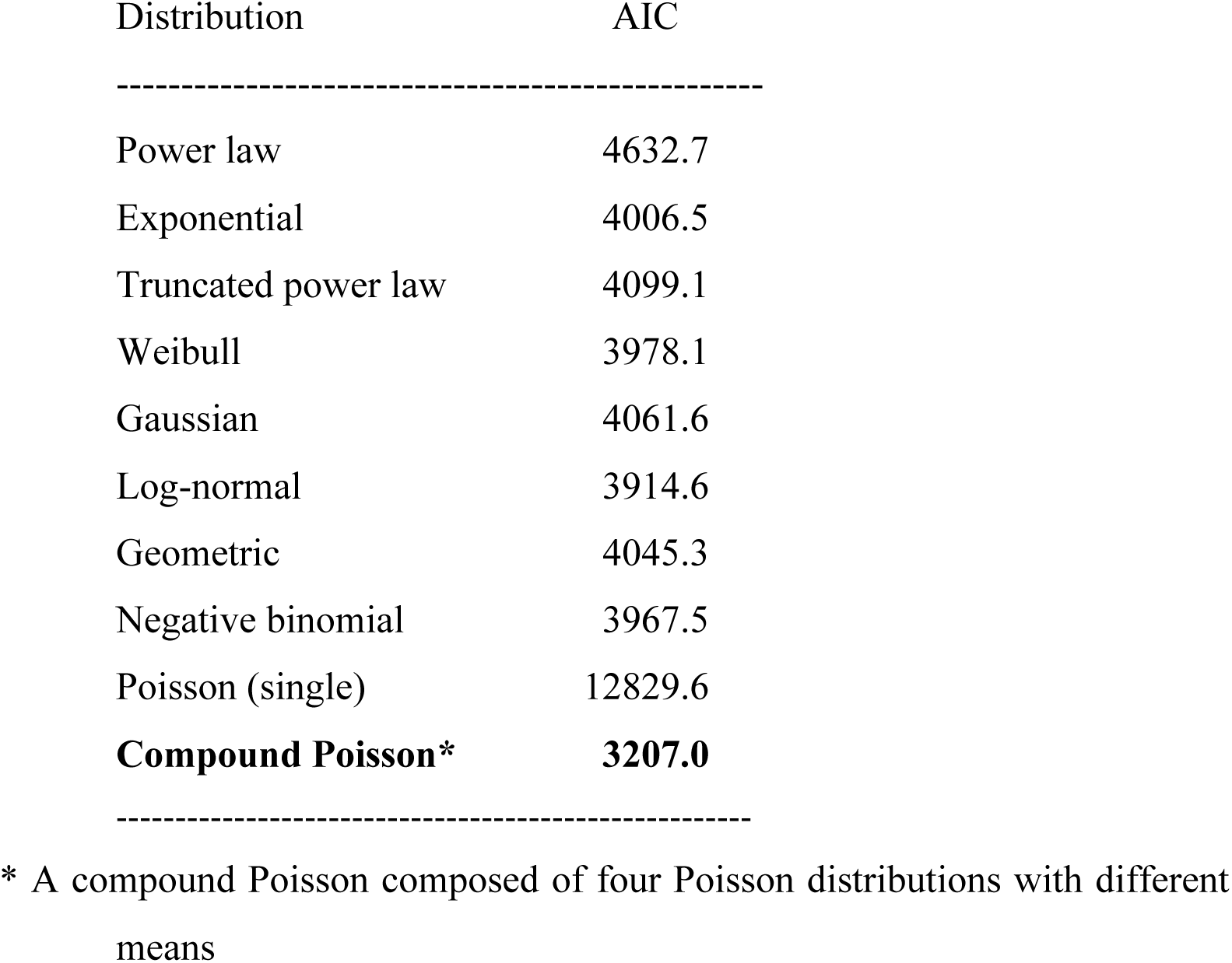
AIC values for the model tested for the distribution of baboon group sizes. The best-fit model (that with the lowest AIC value) is shown in bold.

We also analysed each species separately: Jenks identifies three or four as the optimal number of clusters for all six species, with cluster means that are all close to those found for the sample as a whole and scaling ratios close to 2 (Table 2). More importantly, with the exception of the largest cluster (∼160) for *Papio anubis* and the smallest cluster (∼20) for *P. papio*, *P. hamadryas* and *Theropithecus gelada*, all clusters are present in all species.

**Table 2.** Optimal number of clusters, and the resulting cluster means and mean scaling ratios between successive clusters, for each of the species separately, compared to that for the pooled sample, using the Jenks algorithm.

| Species | Cluster means * |  |  |  |  | Mean scaling ratio |
| --- | --- | --- | --- | --- | --- | --- |
| All *Papio* species | 19.8 (202) | 45.6 (138) | 79.1 (65) | 184.4 (17) |  | 2.12 |
| *P. anubis* | 22.1 (58) | 49.3 (33) | 94.1 (13) | - |  | 2.06 |
| *P. cynocephalus* | 20.7 (32) | 46.6 (41) | 80.0 (28) | 176.0 (4) |  | 2.06 |
| *P. ursinus* | 17.8 (99) | 37.1 (49) | 64.9 (10) | 99.8 (8) |  | 1.79 |
| *P. papio* | - | 42.0 (19) | 113.0 (9) | 216.3 (6) |  | 2.30 |
| *P. hamadryas* | - | 52.7 (7) | 88.0 (3) | 190.0<sup>§</sup> (3) |  | 1.91 |
| *Theropithecus gelada* | - | 54.8 (10) | 131.0 (8) | 237.3 (4) |  | 2.11 |

\* Numbers in parentheses in the body of the table are the number of groups assigned to the cluster by the algorithm.
§ Jenks gives four clusters, but the two largest contain only one and two groups, respectively, with cluster means of 150 and 210. The weighted average is 190.

We suggest that these cluster values make up a set of linear oscillators, with attractors at ∼20 and ∼40, ∼40 and ∼80, and ∼80 and ∼160 respectively. In effect, group size oscillates between the limits set by a given oscillator (i.e. 20/40, 40/80 or 80/160). Within an oscillator, natural growth rates cause groups to increase in size until they fission at the upper value to return to the lower value, and begin the cycle all over again.

It seems that most populations have a characteristic group size that sits comfortably within one of these oscillators. Analysis of the distribution of group sizes for the 27 sites with at least five censused groups indicates that only five are not (unimodal) normally distributed. In these five cases (Amboseli NP, Gilgil, Mt Assirik, Nairobi NP and Giants Castle), the bimodality is created by a small number of very large (N>100) outliers, with the bulk of the groups normally distributed within one or other of the two smaller oscillators. This may reflect the fact that these sites are typically associated with a mosaic of habitat types. The rest all have a group size distribution that falls mainly into one of the oscillators. Fig. 3 plots the proportion of groups in the smallest (20/40) oscillator in each of these populations. The data exhibit a clear bimodal pattern: about half the populations have all or most their groups in the 20/40 oscillator and half have all or most of their groups within the 40/80 or 80/120 oscillators. There is a conspicuous absence of groups with an even split.

**Figure 3.**
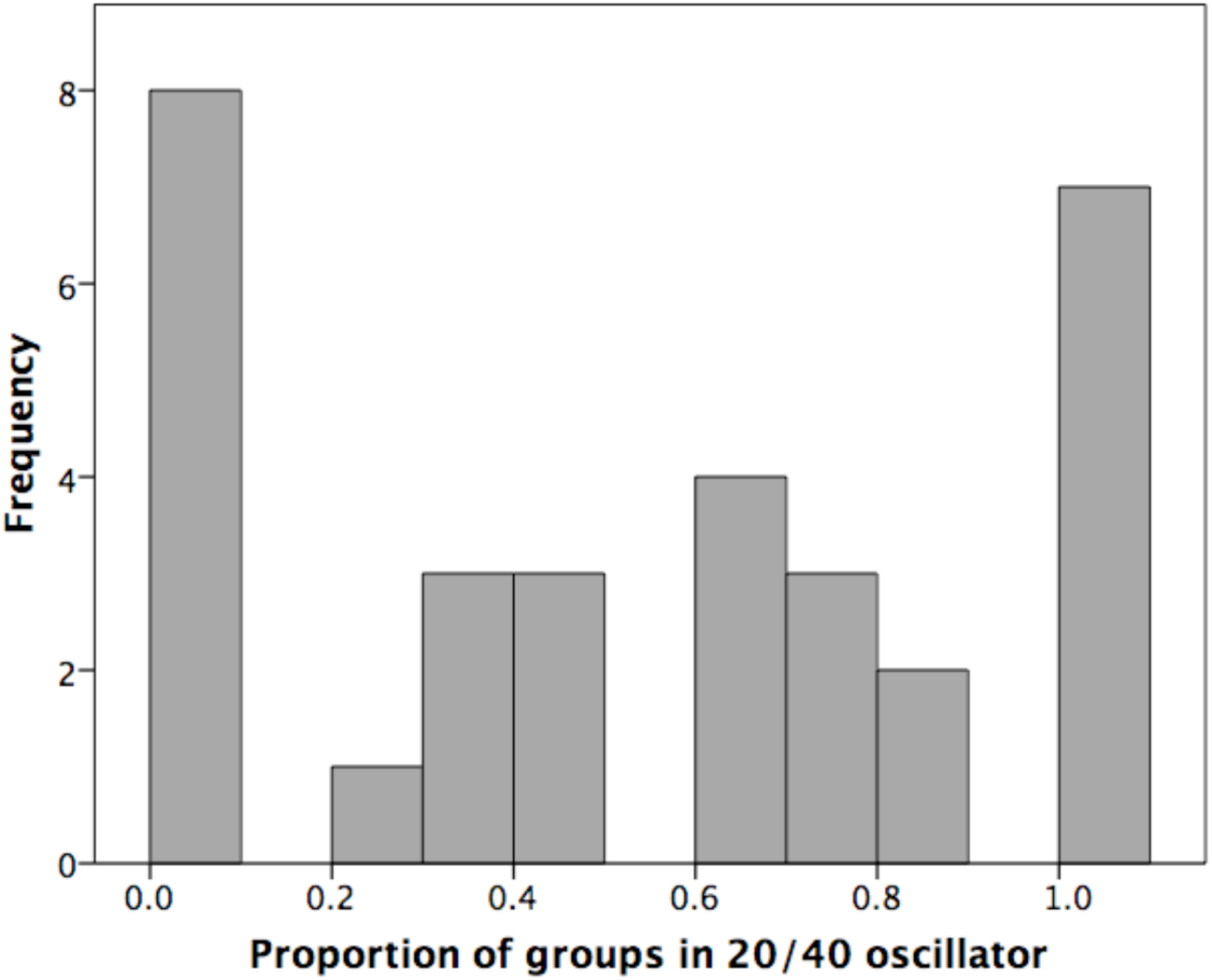
Proportion of groups in populations with >4 censused groups that fell into the lowest oscillator (20/40) for all six species.

This raises three questions: (1) Why are there distinct oscillators rather than a continuous series of optimal values tracking some environmental variable like rainfall? (2) Why do different populations seem to opt for different oscillators? (3) What are the structural consequences of the oscillators? We suggest that the trade off between fertility and predation risk is the key to answering these questions.

### Optimising group size in different habitats

Fig. 4 plots population mean group size as a function of predator density and tree cover (the two main factors that influence predation risk for baboons: Cowlishaw 1997a,b). Predator density is the sum of the densities of the baboons’ two main predators (leopard and lion: Cowlishaw 1994), each predicted from species-specific climate envelope models using data from 59 locations distributed across sub-Saharan Africa (Bettridge et al. 2010). Tree cover (percent of ground covered by tree canopy) is estimated from satellite imagery for the *Papio* populations (see Bettridge et al. 2010) and from ground transects for the gelada populations (Iwamoto & Dunbar 1983). Tree cover is significantly lower in hamadryas and gelada habitats than the habitats of the other *Papio* species (Fig. S1). Essentially, group size is low when there are plenty of trees available as refuges, but, in habitats with few trees, groups are large (in some cases, very large) *if* predator density is high. A general linear model indicates that, while there is no main effect for predation in the predicted direction (F_1,20_=0.75, p=0.199), mean group size is significantly higher in less forested habitats (F_1,20_=3.77, p=0.033) and there is a significant forest*predator interaction (F_1,20_=3.52, p=0.038). (We here test a set of explicit directional hypotheses based on Cowlishaw [1997a,b], so all statistical tests are necessarily 1-tailed.)

**Figure 4.**
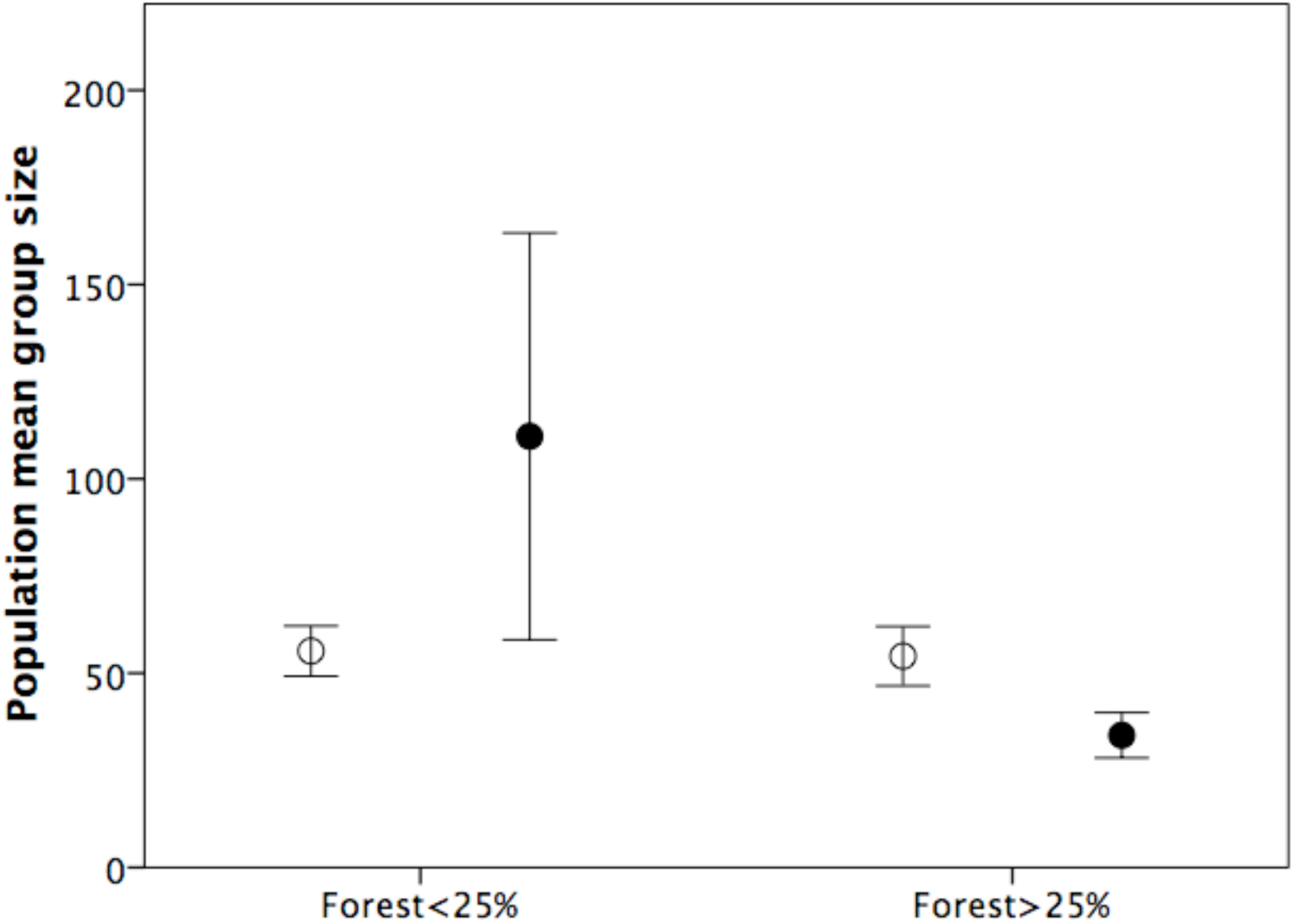
Mean (±1se) population mean group size as a function of predator (lion + leopard) density (filled symbols: >0.25 /km^2^; unfilled symbols: <0.25/km^2^) and percentage tree cover. Data: *ESM* Table S2.

To further illustrate this, Fig. 5 plots mean population group size against a composite index of predation risk. The predation risk index is the sum of the standardised deviate of predator density and the standardised deviate of habitat openness (defined as 100 minus percent tree cover). There is a significant relationship between mean group size and the composite index of predation risk (r=0.506, N=24, p=0.006 1-tailed). In other words, resort to large group sizes (and hence the higher oscillators) appears to be driven by high predation risk.

**Figure 5.**
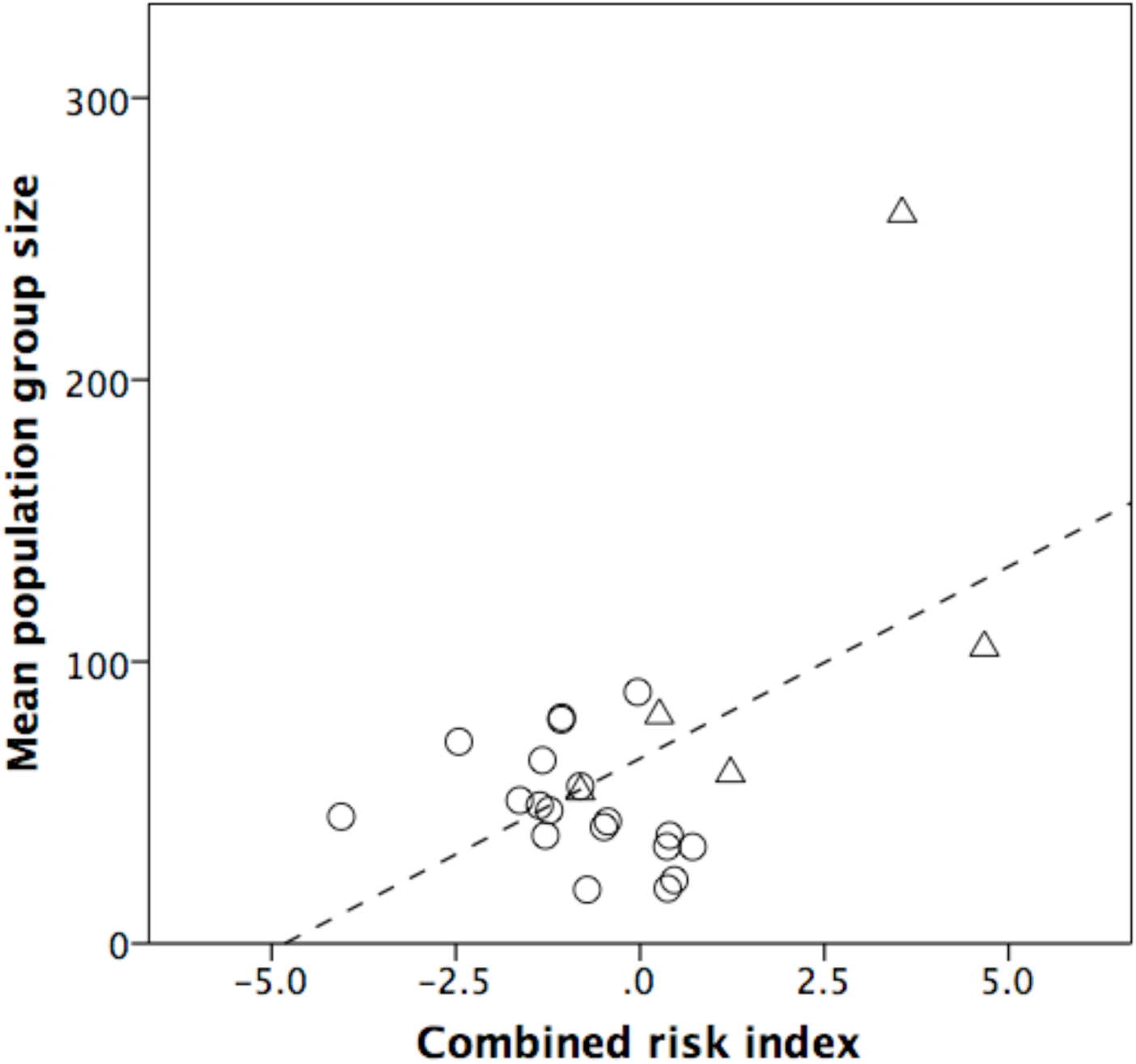
Mean population group size plotted against a composite index of predation risk (standard deviation of predator density plus standard deviation of converse of tree cover [i.e. 100 - % tree cover]). Unfilled symbols: woodland baboon populations; filled symbols: hamadryas and gelada populations. Data: Table S2.

Finally, we use Bettridge et al.’s (2010) baboon time budget model to calculate minimum and maximum ecologically tolerable group sizes as a function of climatic conditions. Minimum group size is a simple function of the density of baboons’ two main predators (leopard and lion) (Bettridge et al. 2010). Maximum group size is determined by first calculating the amount of time that animals need to devote to feeding, moving and resting using the equations given by Bettridge et al. (2010) that relate each of these to climate variables; we then subtract the sum of these from 100% (i.e. all daytime) to give the maximum amount of time available for social grooming; finally, we use the regression equation relating required grooming time to group size in primates (Dunbar 1991; Lehmann et al. 2007a) to determine the largest group size that the animals’ time budget could support at that locality (for details, see Dunbar et al. 2009; Bettridge et al. 2010).

To simplify the presentation of these results, we transformed the eight climate variables used in the model into functions of a single variable (mean annual rainfall) (using the data given in Bettridge et al. 2010; linear regressions: 0.366≤r≤0.892, p≤0.078 2-tailed) and use these to plot minimum and maximum group sizes across the full range of rainfall regimes (0-2000 mm per annum) found in Africa (Fig. 6). Rainfall is a reliable index of forest cover (tree density) in Africa. Minimum group size declines slowly as rainfall increases (i.e. as tree density increases), with a mean value of ∼20.7 (range 18.6-25.9). Maximum tolerable group size increases with rainfall, rising from a base of ∼55 to a maximum of ∼110 in the wettest (i.e. richest) habitats. Maximum group size sets an absolute upper limit and groups cannot exceed this value without paying a price in terms of group cohesion or energy intake because they will not be able to balance their time (and hence energy/nutrient) budget. The space between the minimum and maximum lines in Fig. 6 represents the genus’s realisable niche space: groups will be in ecological balance only if they lie between the two lines (Dunbar et al. 2009). Although groups can stray outside these limits, they should be able to do so only to a limited extent and, usually, only for a limited period of time.

**Figure 6.**
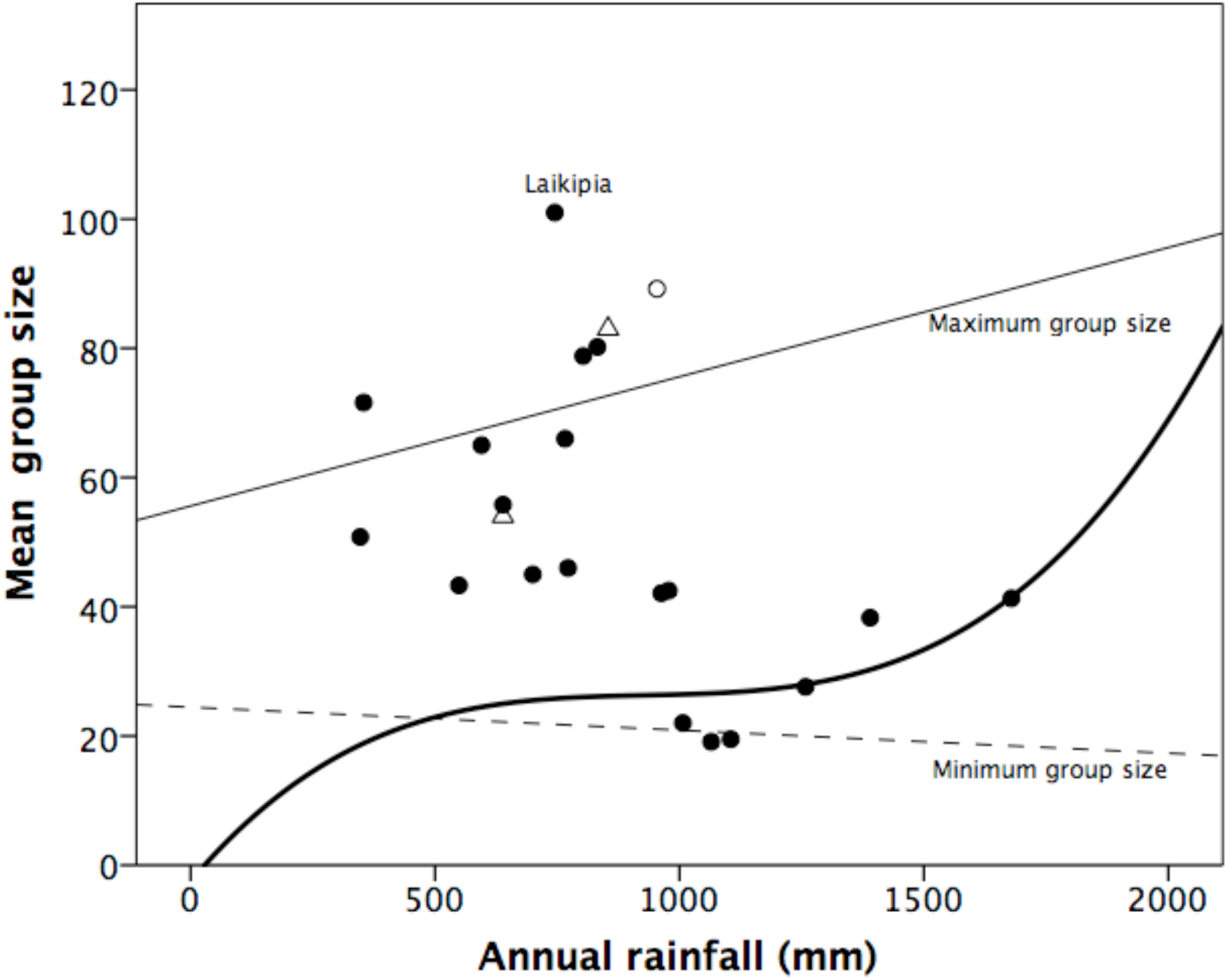
Mean group size for individual baboon populations, plotted against annual rainfall. Filled circles: woodland baboon populations; open circle: Guinea baboon population; triangles: hamadryas populations. The solid line gives the maximum ecologically tolerable group size (*N_max_*) determined by the time budget model, and the dashed line gives the minimum group size (*N_min_*) determined by predation risk, using the equations from Bettridge et al. (2010) and converting all climatic variables into functions of annual rainfall (using regression equations based on data given by Bettridge et al. 2010). Heavy solid line plots percentage of tree cover, based on data given for baboon study sites by Bettridge et al. (2010). See text for details.

Notice that the lines on Fig. 6 represent the two conventional components of fitness (fertility and survivorship). In any given habitat, animals will reduce predation risk, and hence maximise survivorship (albeit at the expense of fertility), by living in the largest groups they can (i.e. closest to the upper dashed line); conversely, they will maximise fertility (at the expense of increased predation risk, and hence reduced survivorship) by living in the smallest groups they can (i.e. closest to the lower solid line). Where they choose to position themselves in this state space should depend on how intrusive predation risk is for them.

The mean group sizes for 19 populations of *Papio* for which detailed climate data are available are plotted onto Fig. 6 as a function of their actual local rainfall regime. All except one of the datapoints lie within or close to the minimum and maximum lines. The exception (Laikipia, Kenya) which lies just outside the upper limit is a site with a particularly low density of tree cover (10%) and a high predicted density of predators (and, within the last century, an unusually high actual density of leopard). More importantly, there appears to be a phase shift in group size at ∼1000mm rainfall: populations seem to switch from maximising group size in dry habitats to minimising group size in wet habitats. This likely reflects the availability of trees that can act as refuges when a group encounters predators during foraging: the heavy sigmoid line plots the percentage of tree cover as a function of rainfall. The switch point seems to correspond to a phase transition in the degree of forest cover, with a value of ∼30% tree cover (i.e. quite heavily wooded habitats) demarcating this transition point. In more open habitats, baboons rely on maximising group size to deter predators (at the expense of reduced birth rates), whereas more wooded habitats allow them to live in smaller groups so as to maximise fertility.

### Fertility

Next, we ask how fertility is affected by group size. Fig. 7 plots mean birth rate against group size for 25 baboon study sites (including three for gelada). The woodland baboon data are clearly best explained by a quadratic regression (b=0.149+0.014N-0.00016N^2^, where b = annual birth rate per female and N = group size; F_2,12_=6.82, r^2^=0.532, p=0.011; linear: F_1,13_=0.30, r^2^=0.022, p=0.594). The quadratic relationship holds individually across all three woodland species (Dunbar et al. 2018a), and when the Guinea baboon datapoints are included (F_2,14_=5.38, r^2^=0.434, p=0.019). It seems that fertility increases monotonically with group size to a maximum of about 0.54 births/year at a group size of ∼55, and then declines again at a similar rate.

**Figure 7.**
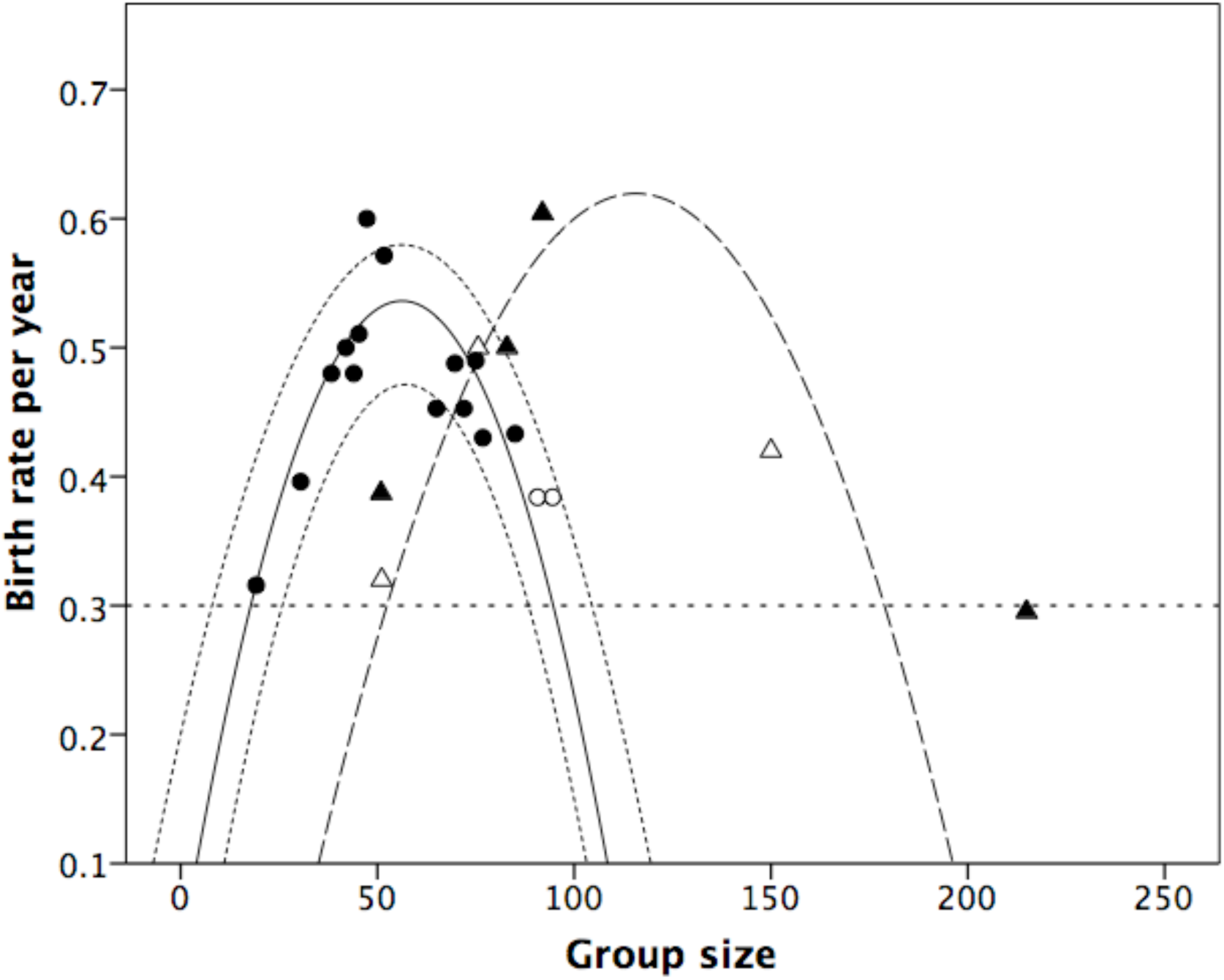
Mean fertility (births per adult female per year) for individual baboon groups, plotted against group size. Filled circles: woodland baboons; open circles: Guinea baboons; solid triangles: hamadryas; open triangles: gelada. The best fit regression for the three woodland species combined has a quadratic form (solid line, with 95% CI indicated by dotted lines). A separate regression (dashed line) is set through the hamadryas and gelada datapoints, with a quadratic again being the best fit.

Hill et al. (2000) found that *Papio* birth rates are also independently determined by mean ambient temperature (an index of habitat quality: Dunbar et al. 2009). However, for the woodland baboons at least, the results are exactly the same if we plot residuals from a regression of fertility on rainfall against group size (Fig. S2), indicating that, irrespective of the effect that habitat quality has on birth rates, birth rates are independently a quadratic function of group size.

In contrast, the hamadryas and gelada datapoints lie well outside the 95% confidence intervals for the woodland species. Their data also fit a quadratic regression (F_2,8_=10.66, r^2^=0.727, p=0.006; linear: F_1,19_=1.09, r^2^=0108, p=0.323) that is very similar in form, but with a significantly lower intercept (meaning that the graph is displaced to the right compared to that for the woodland baboons). For the hamadryas and gelada, a quadratic relationship gives a maximum birth rate at groups of N≈125 (i.e. slightly more than twice that for the woodland species). This result holds even when we use the maximum group sizes for hamadryas (sleeping troops) and gelada (the band) (Fig. S3).

Fig. 7 indicates that no population has a birth rate below 0.3 per annum. With a ∼13-year reproductive lifespan, a minimum fertility of ∼0.15 births per year is required for replacement (i.e. two offspring at the end of a lifetime). Since survivorship to puberty averages ∼50% in wild baboons (Altmann et al. 1977; Sigg et al. 1982), this would require a minimum birth rate of 0.3 for demographic stationarity (i.e. for a population to avoid extinction). This being so, the regression equation for the woodland baboon populations would set an upper limit to group size at 95 for demographic viability. This value includes 92.4% of all baboon groups in the sample in Fig. 1 (or 93.1% of the groups for the three woodland species: Table S1). Note that this is only just above the mean group size (91.6) for Guinea baboons, suggesting that they are on the edge of viability. Groups larger than this will not be reproducing fast enough to compensate for natural mortality, and will therefore oscillate perpetually around this value until they fission. In contrast, the fertility data for hamadryas and gelada predict a limiting group size of ∼180 (2.2 times the equivalent for the other species), just above the uppermost attractor in Fig. 1.

The quadratic form of this equation also implies a minimum limit on group size: in woodland species, groups smaller than ∼15 would have birth rates below the limit for demographic viability, and so would be in terminal decline. Only 8.6% of all groups in the savannah sample are smaller than this. Note that baboons thus have a lower limit on group size that is substantially larger than the size of groups characteristic of most primate species (∼60% of primate species have a mean group size of <15 individuals: Dunbar et al., 2018b). In contrast, hamadryas and gelada would have a lower limit on group size of 38. The smallest hamadryas band recorded in the present sample is in fact 38 (Table S1), while the mean size for gelada teams (the natural grouping for this species, and the set of harems that stay together most closely) is 34 (Mac Carron & Dunbar 2016).

## Discussion

The regular patterning in the distribution of baboon group sizes suggests a fractal sequence with attractors at ∼20, ∼40, ∼80 and ∼160 that are particularly common. Since these values have a scaling ratio of ∼2.0, they are likely to be the product of a binary fission process, as also seems to be the case in the gelada (Mac Carron & Dunbar 2016). At the same time, very small groups of the size common in many other primates (<15 members) are rare (Fig. 1). In fact, 60% of groups with N<15 in the baboon sample were from South African *Papio ursinus* populations living in high latitude, high altitude (i.e. poor quality, thermally challenging) habitats with extremely low predator densities and environmental conditions that impose severe limits on group size. There are no census data available for *Papio kindae*, but the count of 83 for one group given by Weyher et al. (2014) is within the range for the middle oscillator and similar to the group sizes reported from *Papio cynocephalus* populations. Since minimum group size is determined by female body mass in addition to predation risk (smaller species live in bigger groups: Bettridge & Dunbar 2012), we would expect *P. kindae* to live in larger groups than any of the other species in comparable habitats.

If groups of ∼20 are the minimum viable size, then the corresponding minimum size at fission will need to be around 40-45 – or above if groups do not fission into equal halves. While predation risk and fission processes seem to provide a natural explanation for a fractal pattern of group sizes, they do not of themselves explain why fission should typically occur at the group sizes where seems to, especially as these are well below the limiting group size for baboons set by their carrying capacity in most habitats (Bettridge et al. 2010).

The impact of social competition on fertility is clearly an issue for group-living primates, as it is for all social mammals. The fact that, in both monkeys and humans, the fertility decline can be attributed directly to the number of females in the group rather than to the number of males or total group size (Dunbar 1980; Hill et al. 2000; Ji et al. 2013), and does so even in captivity where food is abundant (Garcia et al. 2006), points to the importance of a social, rather than ecological, explanation – even though competition for food may sometimes provide the context for this. These costs appear to be so strong that they effectively limit group size to ∼15 individuals in most primates (Dunbar 2018). Cercopithecine primates seem to have evolved female coalitions as a mechanism for mitigating these costs in order to make larger groups possible (Dunbar 2012, 2018; Dunbar & Shultz 2017). Female baboons with larger grooming-based coalitions experience less harassment (Dunbar 1980, 2018), have lower cortisol titres (Crockford et al. 2008; Wittig et al. 2008), produce more offspring and live longer (Silk et al. 2003, 2009, 2010).

We have suggested that woodland baboon species are constrained by the fertility curve in Fig. 7 to groups between ∼20 and ∼80 in size, and that this gives rise to a pair of linear oscillators (20/40 and 40/80). We have shown elsewhere (Dunbar et al. 2018b) that the two lower oscillators (20-40 and 40-80) are an evolutionary stable strategy (ESS) for the woodland baboon species. The estimated lifetime reproductive outputs of the average female in the two oscillators, given the fertility rates in Fig. 7, are equal only when the phase shift between them occurs at a group size of ∼40. Phase transitions set either side of this value will result in one or other of the oscillators being disproportionately favoured: if the switch over occurs at smaller group sizes (groups in the range 25-35), then females in the larger oscillator do significantly better, while the reverse is true if the switch over occurs at larger group sizes (45-75). This seems to be due to the fact that a phase transition at the group size that maximises fertility results in females doing equally well either way, the difference simply being whether they experience maximum fertility early or late in their reproductive lives. In other words, these two oscillators seem to represent an optimal partition for females trying to maximise their fitness under different predation risk conditions. Although predation risk might favour a quantitative response, the fertility costs result in a qualitative switch between states (i.e. oscillators).

That the upper values in each case are fission points is indicated by evidence that *Papio ursinus* groups living in a low predation risk habitat in South Africa typically fission at a size of ∼32 (for a mean population group size of 22.4, N=61) whereas *P. cynocephalus* groups living in a high predation risk habitat in East Africa do so at ∼65 (mean population group size 50.7, N=51) (Henzi et al. 1997). These are likely to be facultative responses to environmental conditions, so populations can be expected to switch between these oscillators as environmental circumstances change. Some evidence for this is provided by the fact that, over a period of four decades, the *Papio anubis* population at Gombe switched from a 20/40 range to a 40/80 range and, subsequently, back again (Tony Collins, pers. comm.), though the circumstances that prevailed on them to do so remain unclear.

We are left with the question of why hamadryas and gelada have opted for an oscillator that lies outside this range. Part of the answer may lie in the fact that these two species live in much more open (hence even more predator risky) habitats than those typically occupied by other baboons (Fig. 5). This does not necessarily mean that they have higher predator densities, but it does mean that the availability of large trees to act as refuges is very limited. It is the virtual absence of large trees that obliges hamadryas to congregate at night on rocky outcrops and gelada to seek refuge on sheer cliff faces.

The problem for both these two species is that they can only increase group size significantly to compensate for the higher predation risk providing they can find some way of circumventing the fertility trap. To illustrate the magnitude of this trap we can use the regression equation for fertility from Fig. 7 to model the change in group size over time for groups with initial sizes of 15, 17, 20 and 25, subject to an annual mortality rate that varies randomly across a normal distribution with a mean of 10% and a range of 4-16% (based on data from 3 long term study sites: Moremi, Amboseli and Gombe; Cheney et al. 2004; Bronikowski et al. 2002). Reproductive females are assumed to constitute ∼30% of the group, as is typical of both *Papio* baboons in particular (Fig. S4) and primates in general (Dunbar et al. 2018a). Note that the point of this model is simply to observe the consequences of the particular demographic configuration implied by the fertility rates in Fig. 3.

Fig. 8 plots the results. Groups whose initial size is <17 animals will always go extinct (grey shaded region) because their fertility rates are too low to offset mortality. Groups with an initial size >17 grow slowly for a number of years and then accelerate until they reach an asymptotic value of ∼100, where they will stay indefinitely. When group size exceeds this value, fertility drops and excess mortality reduces group size; this allows fertility to increase, which in turn allows the group to grow again until fertility once more drops below mortality. Left to their own devices, populations will have natural limits at ∼20 and ∼100, and groups will often be stuck at these values for some considerable time (in the first case, until random fluctuations in mortality and female cohort size allow them to break out of the lower cycle; in the second case, until fission radically reduces group size).

**Figure 8.**
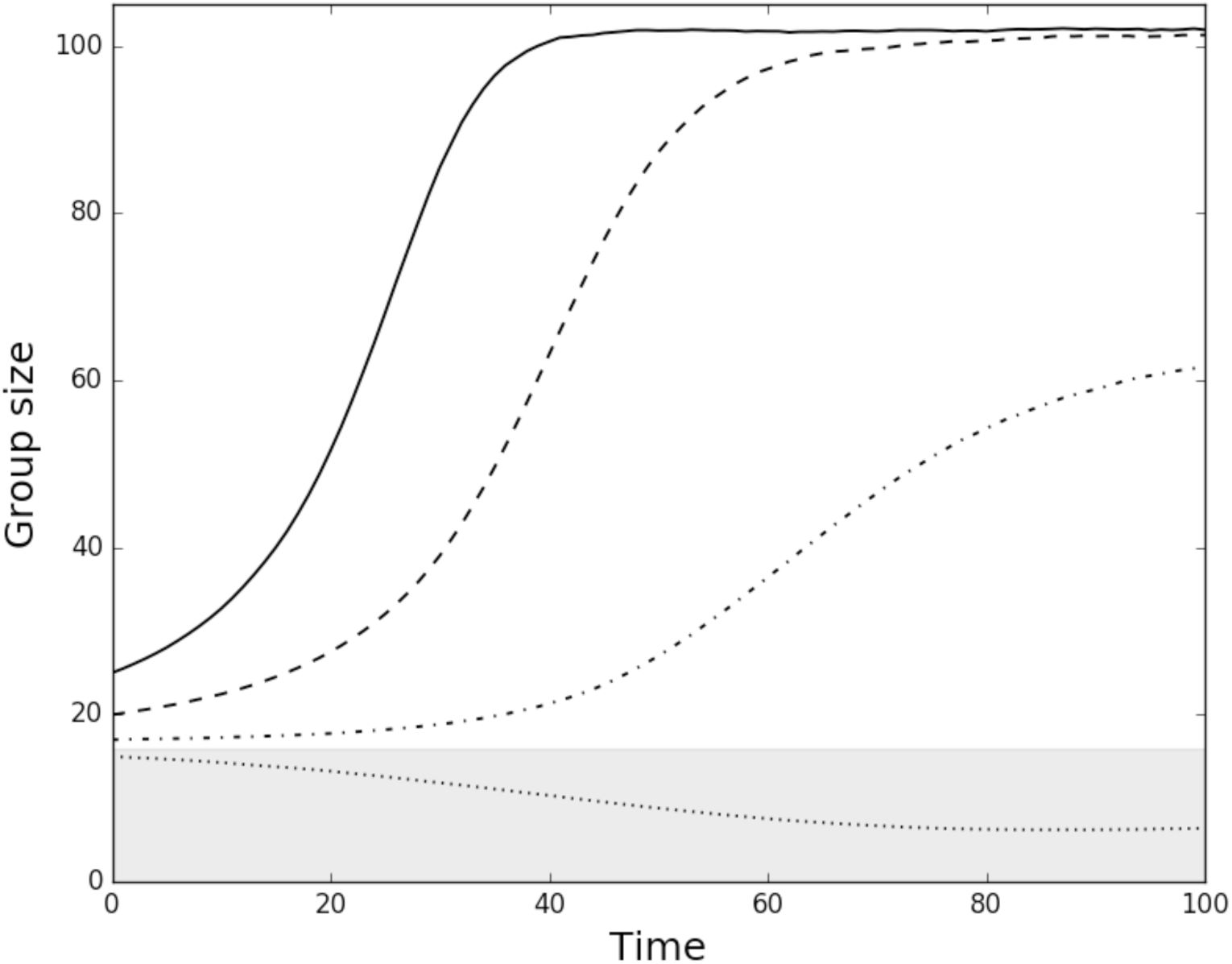
Group size over succeeding time units (years), given the birth rates shown in Fig. 7 for woodland species and an annual mortality rate varying across a normal distribution with a mean of 10% and varying between 4-16%, for different initial group sizes (N=15 = dotted line; N=17 = short dash; N=20 = long dash; N=25 = solid). The lines are mean values for 10,000 simulations in each case. If a group fissions at a size of ∼60, it might do so into daughter groups of 20 and 40 (heavy arrows) or two groups of 30. Grey zone: region (N<17) within which a group will inevitably go extinct because its death rate will always exceed its birth rate.

The contrast between hamadryas and gelada and the woodland species (including the Guinea baboon) is striking. The decline in fertility at group sizes above 50 is sufficiently steep that it will place groups of >100, as occur regularly in hamadryas and gelada, under significant demographic pressure with birth rates below the level for demographic replacement. Fig. 8 suggests that, all else equal, it should be extremely difficult, if not impossible, for these two species to increase group size to the levels that they have been able to do. Yet they have both clearly managed to do so. They have somehow been able to shift their fertility schedule to the right such that they can live in much larger groups (Fig. 7).

Hamadryas live in a poor quality semi-desert habitat, with very few large trees to act as refuges (even along watercourses) and predator densities similar to those faced by woodland baboon populations. In a habitat virtually devoid of large trees, they are obliged to congregate at night in very large herds (typically several bands) on the few rocky outcrops large enough to provide safe night time refuges (Kummer 1968; Sigg & Stolba 1981; Schrier & Swedell 2012b). That predation is a concern for them is indicated by the fact that, during the day, hamadryas bands (which normally disperse over a wide area when foraging) will bunch defensively when predators are heard or sighted (Schrier & Swedell 2012a), just as savannah baboons do (Altmann & Altmann 1970).

Although the gelada live in a cool, high altitude habitat with rich grassy swards that allow very large herds to congregate (at least during the wet season: Mac Carron & Dunbar 2016), they also face a significant level of predation risk (historically from leopard, but more recently hyaena in large numbers) in a completely treeless habitat that can be nutritionally challenging for them during the dry season (Dunbar 1984; Mac Carron & Dunbar 2016). As with the hamadryas, large groups remain a crucial adaptation for minimising predation risk if gelada are to forage on the plateau tops away from the safety provided by the cliff faces of the gorge systems. Where gelada remain on the cliff faces, they have much smaller bands (typically 40-70 animals) and individual harems often forage on their own (Dunbar & Dunbar 1975; Dunbar 1984; Dunbar & Mac Carron 2016).

The dietary adaptations of these two species (Dunbar & Boese 1991) and the dynamics of their social relationships (Dunbar 1983b) are radically different and unlikely to be the cause of this substructuring. The only aspect of behaviour or ecology that is common to the two species is the subdivision of groups (bands) into harems. If so, this provides a principled explanation for why hamadryas and gelada have such different social systems to the other baboons (and the other large-bodied cercopithecines like the mangabeys and macaques). It is perhaps worth noting that when habitat conditions dictate foraging groups significantly smaller than band size in gelada, the harems are sufficiently independent to be able to separate out and forage alone (e.g. Bole valley: Dunbar & Dunbar 1975; Bale Mountains: Mori et al. 1999).

One plausible suggestion, then, is that the hamadryas and gelada have achieved this transition by exploiting a form of alliance-based substructuring (i.e. harems) so as to buffer females against the stresses of living in very large groups. It may be no coincidence that both these species have adopted a harem-based form of sociality in which groups of closely related females (Dunbar 1984; Swedell 2002; Städele et al. 2016) are attached to a male rather than dispersing in small, unstable foraging parties, as happens in chimpanzees (Lehamnn et al. 2007b) and spider monkeys (Korstjens et al. 2006) under similar circumstances. In both hamadryas and gelada, the substructuring is reinforced (more so in the first case, less so in the second) by forms of male herding behaviour that are not observed in the other species, but functionally the consequences are that groups of (usually related) females associate with a male capable of providing additional support during altercations without creating the fertility problems associated with associating with more females. In this respect, they are rather similar to gorilla groups, where the male also provides what amounts to a bodyguard service (Harcourt & Greenberg 2001).

But why should such substructuring take the form of harems associated with a single breeding male (and sometimes an associated follower male)? One possible explanation might be that the size of coalitions is limited by matriline size, and this itself may set a limit on the point at which fertility is maximised by woodland baboons in Fig. 7. Matriline size is likely to be limited by lifehistory processes, since matrilines typically fission once their matriarch dies. If this sets an upper limit to matriline size, then their ability to buffer females against the stresses of group-living will set a corresponding limit to group size. Some evidence to suggest this is given by the fact that, across cercopithecine primates, mean grooming clique size (commonly a matrilineal grouping) correlates with species mean group size (Kudo & Dunbar 2001; Lehmann & Dunbar 2009b). Perhaps the only way of increasing coalition size beyond this may be to add a male who can act as a ‘hired gun’ to protect the females from harassment, but who doesn’t himself compete with the females (the ‘bodyguard hypothesis’: Mesnick 1997; Wilson & Mesnick 1997; Harcourt & Greenberg 2001). The effectiveness of this is evident in gelada. Conflicts between neighbouring harems are usually initiated by one, occasionally two, females; if the conflict escalates, more females will become involved, until eventually the harem males are drawn into the dispute in defence of their females (Dunbar 1983a, 2018).

In many ways, Guinea baboons (*Papio papio*) face similar problems: the density and height of the ground level vegetation throughout most of their range in West Africa, combined with a relatively flat landscape and a low overall density of large trees away from the major rivers, makes their habitats particularly predator-risky. The density and height of the ground-level vegetation, for example, means that Guinea baboons have little advance warning of the approach of cursorial predators, thus making them especially vulnerable (pers. obs.; see also Cowlishaw 1997b). The Guinea baboon social system is not well understood, although most authors agree that some form of loose multi-level grouping seems to be involved. Patzelt et al. (2011) provide some evidence (albeit based only on association patterns between individual males) that this species has a three tier social system that consists of parties, gangs and communities. They identify the gang (defined as the set of animals that share the same home range) of around ∼60 animals (at their study site) as the equivalent of the baboon troop (and hamadryas band), with the party being a group of females loosely associated with a male. The *Papio* time budget model (Bettridge et al. 2010) gives a maximum ecologically tolerable group size of 88.5 for baboons in the habitats occupied by Guinea baboons in southeast Senegal – almost exactly the observed mean group size for the censussed Guinea baboon populations from this area (Fig. 2). This suggests that this species is attempting to maximise its group sizes, presumably to cope with high predation risk, but, in contrast to the hamadryas, does so at the expense of reduced fertility and social coherence because it has not mnaged to evolve the behavioural and cognitive mechanisms (e.g. the male herding behaviour observed in both hamadryas and gelada) that would allow it to substructure these groups effectively. This might explain their seemingly somewhat chaotic social structure.

These analyses provide principled answers to the questions we posed in the light of the fractal distribution of baboon group sizes, namely why there are distinct oscillators rather than a series of optimal values, why different populations prefer different oscillators and what the structural consequences of the oscillators are. It seems that the oscillators represent an evolutionarily stable strategy set driven by local predation risk. While baboon sociality can cope with the two lower oscillators without significant stress, the upper oscillator (characteristic of hamadryas and gelada) is only stable if the animals can introduce structural adaptations that allow the fertility schedule to be shifted sufficiently to the right. This seems to have involved formal substructuring that has involved an intensification of the natural matrilineal coalitions characteristic of baboons and other cercopithecines and then attaching these subgroups to a male who acts as a ‘hired gun’. At least in the case of the hamadryas, this seems to have required some degree of genetic underpinning in terms of male herding behaviour (Nagel 1973; Bergman et al. 2008), a pattern that seems crucial in enforcing their harem-based system (given that both anubis and hamadryas females seem able to adapt to either social system: Kummer 1968). However, even gelada males herd their females away from rivals (Dunbar 1984), although they do not do so as effectively as hamadryas males do.

There are several predictions that follow from this model of baboon demography. First, the oscillator should change when baboons migrate into a more, or less, predator risky habitat, or there are changes in, for example, tree density due to deforestation or longterm climatic change influencing habitat structure and tree composition (as has happened at Amboseli: Altmann et al. 1985). Second, individual groups should show cyclic changes in average female fertility over time, following the patterns in Fig. 7. These changes should be upwards (followed by rapid collapse) for baboon groups in the 20-40 oscillator, and downwards (followed by rapid improvement) in populations in the 40-80 oscillator.

## Acknowledgments

This research has been funded by a European Research Council Advanced Investigator grant (#295663) to RD. We thank Cole Robertson for collating the *Papio* database.

## Supplementary Information

### Supplementary Methods

We take group (or troop) size to be that defined by the field worker: in most cases, this grouping is fairly obvious, since the animals forage and sleep together, and maintain a degree of demographic coherence and stability as well as spatial separation from other similar groups. For *Papio hamadryas*, we follow convention and take the band to be the homologue of the *Papio* troop – a set of animals that share the same home range (Kummer 1968). Exactly what counts as a group in *Papio papio* has been the subject of debate because of this species’ rather flexible form of sociality (Dunbar & Nathan 1972; Boese 1975; Sharman 1981), but all field workers on this species agree on the existence of some form of stable social group, at least as defined by animals that share a common range area (Patzelt et al. 2011). The data are provided in the *SI* Table S1.

In most cases, fertility rates are based on observed mean birth rate or mean interbirth interval. In a few cases, birth rates were estimated from the number of immatures per female in a group. Immatures are defined as animals that are pre-puberty, with puberty occurring at around 4 years of age (Altmann et al. 1977). In the case of the gelada, we have used mean herd size rather than band size as the appropriate social grouping for comparison with the other *Papio* species: this is because gelada bands are probably not the homologue of baboon troops (MacCarron & Dunbar 2016) but rather are loose clusterings of harems that share a common range area. Not all the units of a band are found together on any given day (herd sizes vary from a single harem to upwards of 50 harems from several bands). Given that the driver of functional infertility is the number of individuals who are foraging or resting together at any given time, mean herd size seems the most appropriate measure.

### Supplementary Results

Fig. S1 plots the distribution of forest cover at sites listed by Bettridge et al. (2010) where the six species have been studied in detail, with gelada habitats from Iwamoto & Dunbar (1983). For *Papio* species, forest cover is estimated from satellite imagery; that those for the gelada sites are based on ground transects carried out by the authors. Hamadryas and gelada live in habitats with especially low levels of forest cover. Note, however, that these data underestimate the contrast between hamadryas and gelada habitats and those occupied by other baboon species: the size of trees is much smaller in most hamadryas and gelada habitats than those in the habitats occupied by the other woodland baboon species. In gelada habitats, for example, few trees are over 5m in height.

**Fig. S1.**
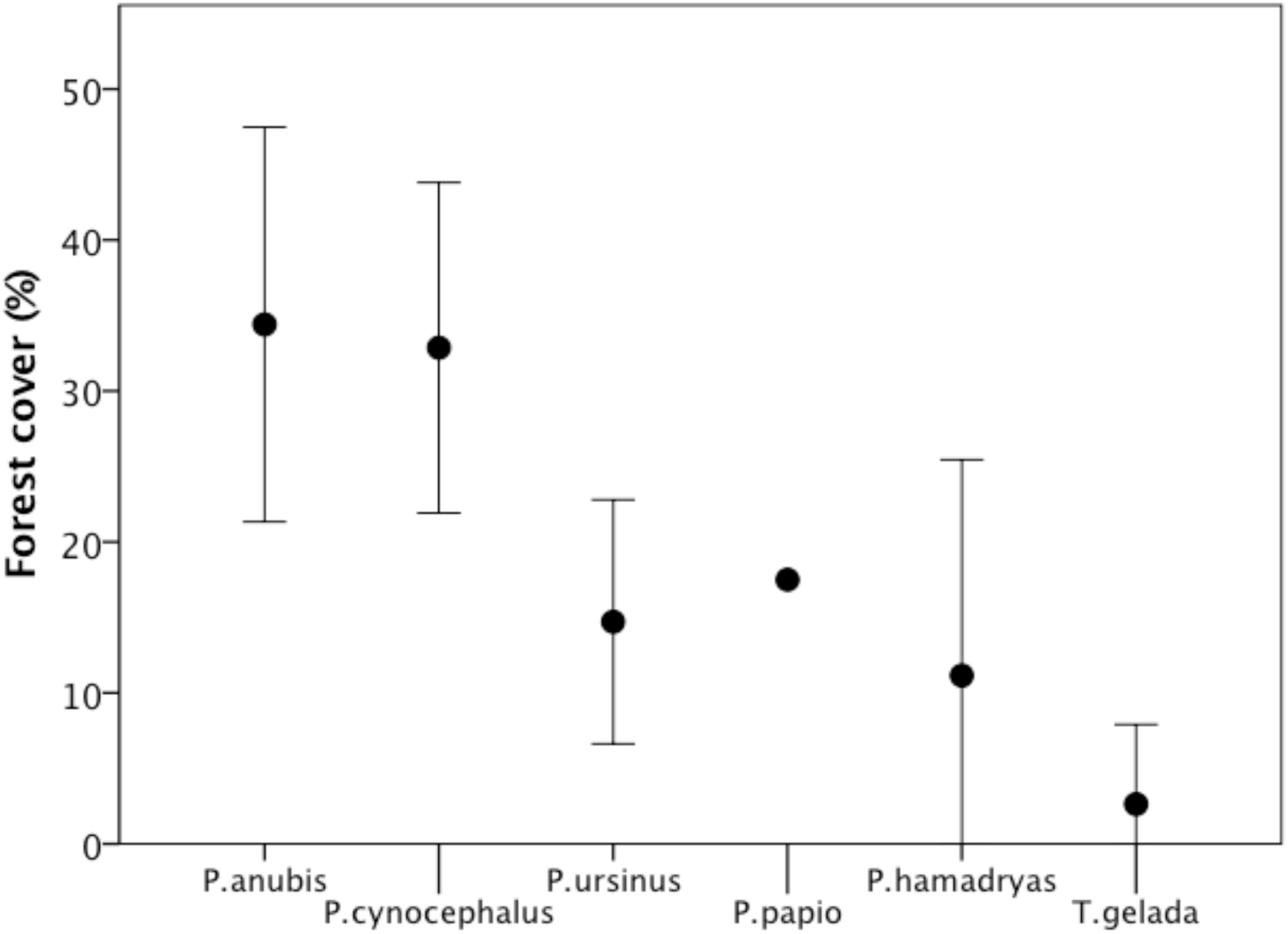
Mean (±2 se) forest cover (indexed as percentage of ground cover) for the six species. Sources: Bettridge et al. (2010) and Iwamoto & Dunbar (1983)

Baboon fertility is independently predicted by both the number of adult females in the group and the mean ambient temperature of the habitat (an index of habitat quality) (Hill et al. 2000). To check whether environmental conditions might be a confound in our results, we regressed birth rate on mean habitat ambient temperature as a quadratic relationship (b = −0.672 + 0.104Temp - 0.00231*Temp^2^: F_2,18_=3.77, r^2^=0.296, p=0.043; linear: F_1,19_=0.01, r^2^=0.000, p=0.933), and calculated residual birth rates from this regression. The results (Fig. S2) are identical to those shown in Fig. 3. Gelada were not included in this analysis because they inhabit a different (very high altitude) temperature regime to *Papio* baboons.

**Figure S2.**
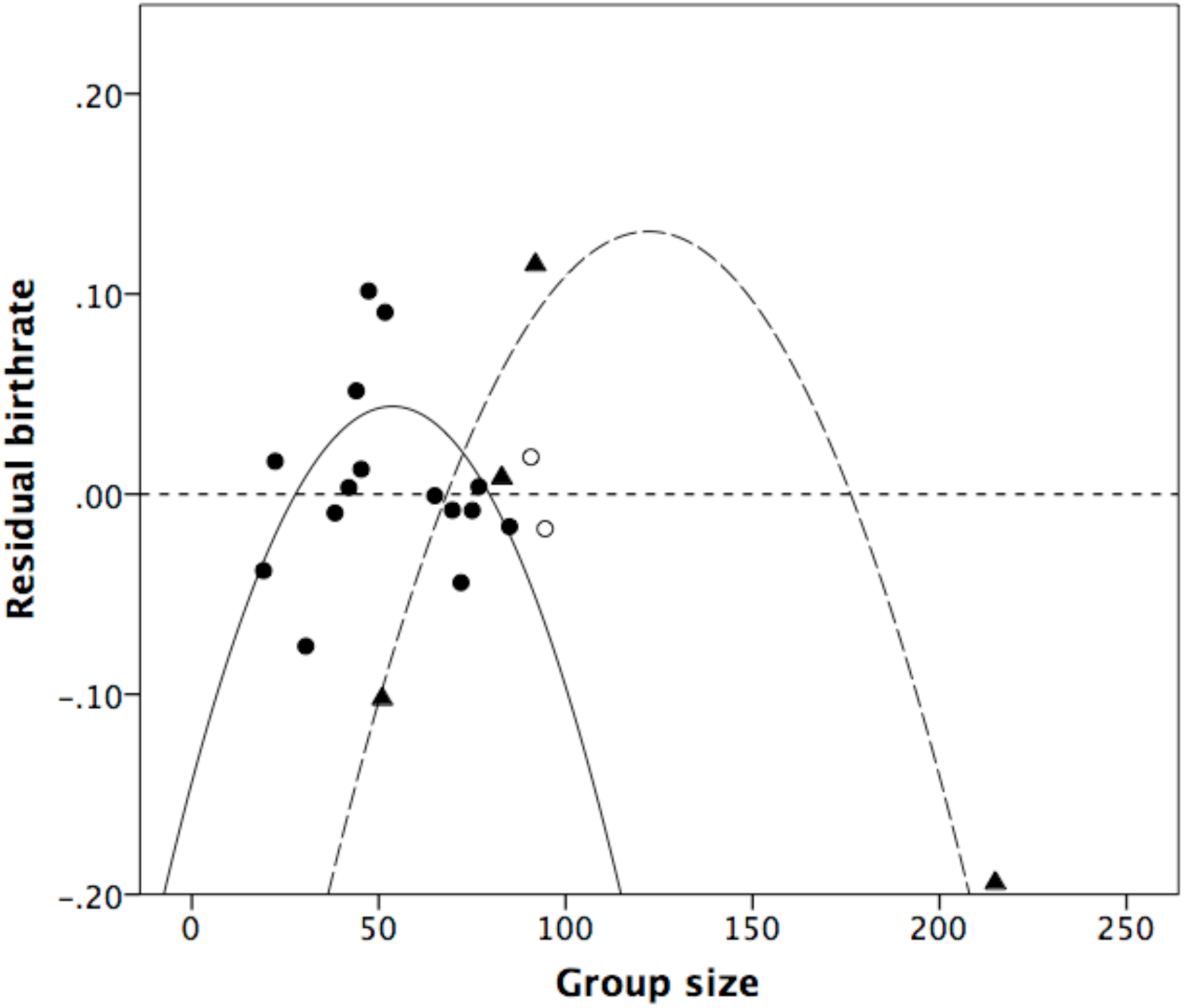
Residual of mean birth rate regressed on mean local temperature plotted against group size for individual populations. Filled circles: savannah baboons (P. anubis, P. cynocephalus and P. ursinus); unfilled circles: P. papio; triangles: P. hamadryas. Separate quadratic regressions set to data for savannah baboons and P. hamadryas, respectively. Gelada are not included (see text).

Fig. S3 confirms that the results of Fig. 3 are invariant with social scale: the same U-shaped pattern emerges when birth rate is plotted against the largest social groupings observed in the two species that live in multi-level social groupings (hamadryas baboons and gelada). The fit, however, is better for Fig. 3, which uses foraging group size (bands for hamadryas, mean herd size for gelada), the groupings in which the animals spend most of their time (and hence where stress effects are likely to be most intense).

Overall, the average number of females in baboon groups is 14.5±6.9, and average group size is 50.3±21.4 (N=16), for a ratio of 0.288. Fig. S5 plots the data for these groups: the regression line has a slope of ∼3. The mean ratio of females to total group size for these individual groups is 0.282±0.05. In fact, this value seems to be characteristic across primates as a whole (data in Campbell et al. 2007).

**Figure S4.**
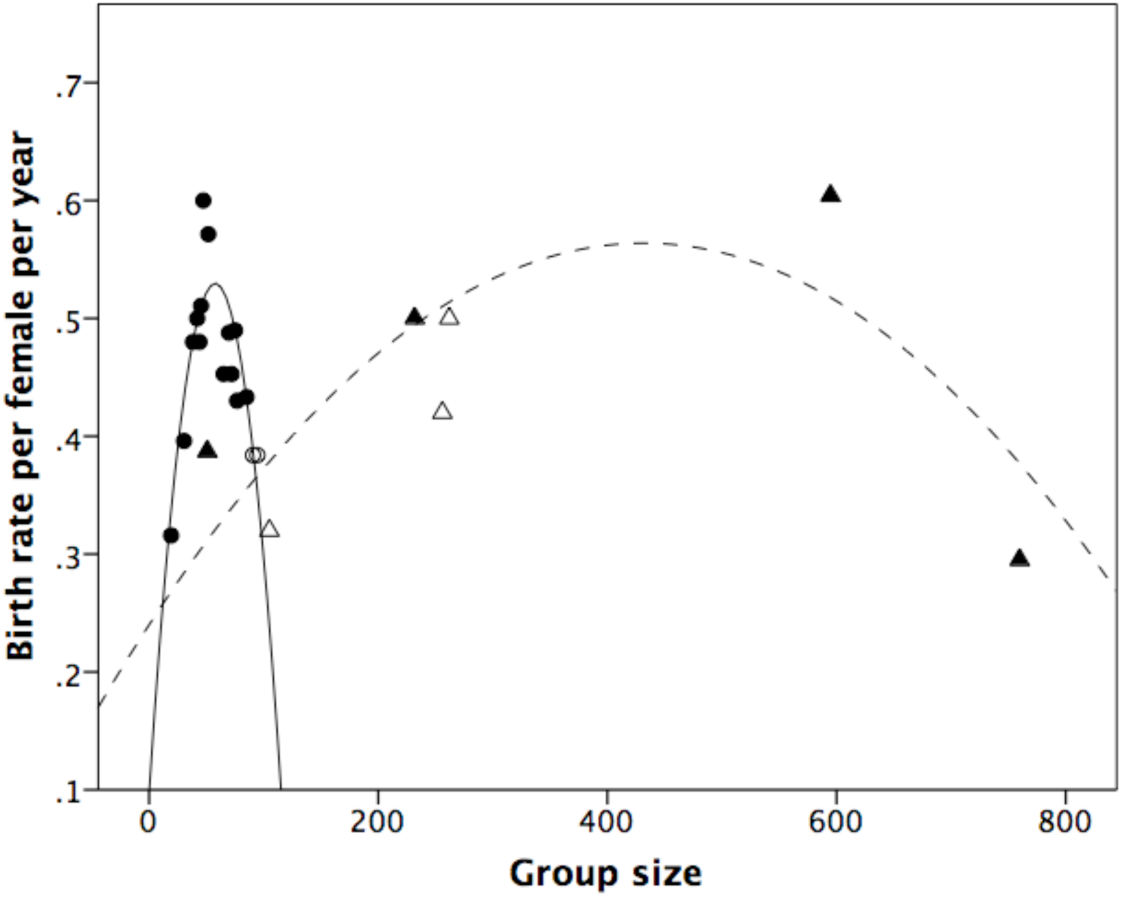
Birth rate plotted against highest level of social grouping (multi-band sleeping troop in the case of Papio hamadryas and band in the case of Theropithecus gelada. Filled circles: savannah baboons (P. anubis, P. cynocephalus and P. ursinus); unfilled circles: P. papio; solid triangles: P. hamadryas. Separate quadratic regressions set to data for savannah baboons and P. hamadryas, respectively.

**Figure S5.**
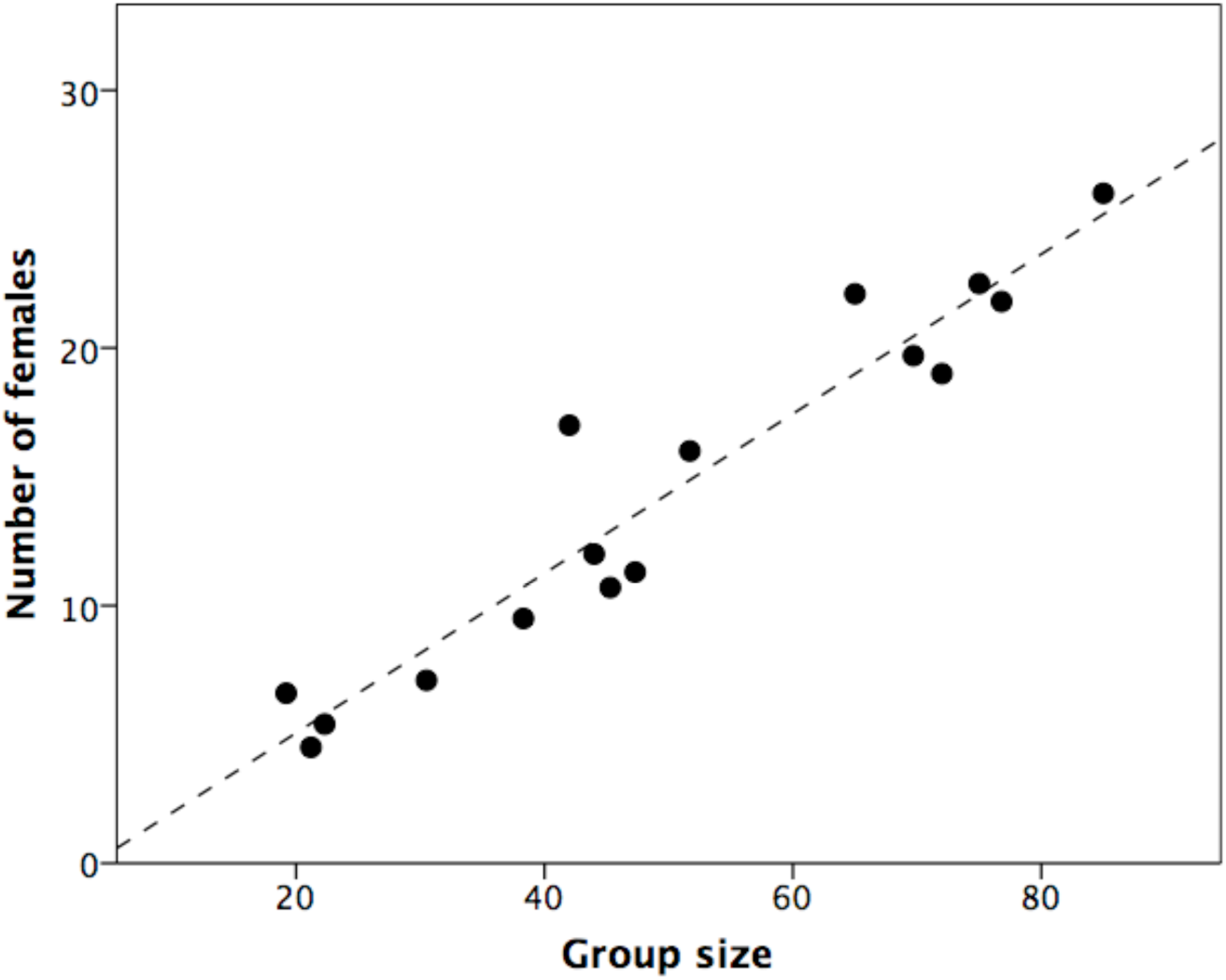
Number of females plotted against total group size for all the savannah baboon groups in the fertility sample. The dashed line is the best fit linear regression and has a slope of ∼0.3.

